# Cytokinin fluoroprobe and receptor CRE1/AHK4 localize to both plasma membrane and endoplasmic reticulum

**DOI:** 10.1101/744177

**Authors:** Karolina Kubiasová, Juan Carlos Montesinos, Olga Šamajová, Jaroslav Nisler, Václav Mik, Lucie Plíhalová, Ondřej Novák, Peter Marhavý, David Zalabák, Karel Berka, Karel Doležal, Petr Galuszka, Jozef Šamaj, Miroslav Strnad, Eva Benková, Ondřej Plíhal, Lukáš Spíchal

**Author notes:** deceased (Petr Galuszka). These authors contributed equally (Karolina Kubiasová, Juan Carlos Montesinos).

## Abstract

The plant hormone cytokinin regulates various cell and developmental processes, including cell division and differentiation, embryogenesis, activity of shoot and root apical meristems, formation of shoot and root lateral organs and others ^1^. Cytokinins are perceived by a subfamily of sensor histidine kinases (HKs), which via a two-component phosphorelay cascade activate transcriptional responses in the nucleus. Based on the subcellular localization of cytokinin receptors in various transient expression systems, such as tobacco leaf epidermal cells, and membrane fractionation experiments of Arabidopsis and maize, the endoplasmic reticulum (ER) membrane has been proposed as a principal hormone perception site ^2–4^. Intriguingly, recent study of the cytokinin transporter PUP14 has pointed out that the plasma membrane (PM)-mediated signalling might play an important role in establishment of cytokinin response gradients in various plant organs ^5^. However, localization of cytokinin HK receptors to the PM, although initially suggested ^6^, remains ambiguous. Here, by monitoring subcellular localizations of the fluorescently labelled natural cytokinin probe iP-NBD ^7^ and the cytokinin receptor ARABIDOPSIS HISTIDINE KINASE 4 (CRE1/AHK4) fused to GFP reporter, we show that pools of the ER-located cytokinin fluoroprobes and receptors can enter the secretory pathway and reach the PM. We demonstrate that in cells of the root apical meristem, CRE1/AHK4 localizes to the PM and the cell plate of dividing meristematic cells. Brefeldin A (BFA) experiments revealed vesicular recycling of the receptor and its accumulation in BFA compartments. Our results provide a new perspective on cytokinin signalling and the possibility of multiple sites of perception at PM and ER, which may determine specific outputs of cytokinin signalling.

---

Fluorescently labelled analogues of phytohormones including auxin, gibberellin, brassinosteroid and strigolactone have been successfully used to map the intracellular fate of their receptors *in planta* ^8^. To adopt this tool for mapping subcellular localization of cytokinin receptors, using docking experiments and cytokinin activity screening bioassays, we selected a fluorescently labelled bioactive compound that interacts with the binding site of a cytokinin receptor. To minimize the possibility of metabolic conversion by cytokinin deactivation enzymes *in planta*, isopentenyladenine (iP) was used as a natural cytokinin that cannot be transformed through *O*-glycosylation at the cytokinin side chain. Furthermore, attachment of 7-nitro-2,1,3-benzoxadiazole (NBD), a small fluorophore, to the *N*9 position of iP eliminates a risk of a metabolic conversion of the final cytokinin fluorescent probe iP-NBD (Fig. 1a) through *N*-glycosylation, or formation of cytokinin nucleotides. Docking simulations using the CRE1/AHK4-iP crystal structure ^9^ suggested that iP-NBD may be fully embedded into the active sites of AHK receptors (Fig. 1b) with micromolar range affinity. The affinity of iP-NBD to cytokinin receptors was measured using bacterially expressed recombinant AHK3 and CRE1/AHK4 that are known to differ in affinity to iP ^10, 11^. Competitive binding assays with *E. coli* expressing either AHK3 or CRE1/AHK4 ^12^ showed that iP-NBD competes for receptor binding with radiolabelled *trans*-zeatin (*t*Z), with K_i_ ~ 37 μM in AHK3 and K_i_ ~ 1.4 μM in CRE1/AHK4 assays, respectively (Fig. 1c). As shown, iP-NBD had lower affinity to AHK3 than to CRE1/AHK4, making this fluoroprobe more specific to CRE1/AHK4. Despite iP-NBD accommodated into the cytokinin-binding pockets of the receptors, it failed to trigger cytokinin response in *E. coli* (*ΔrcsC, cps∷lacZ*) receptor activation assay with recombinant AHK3 and CRE1/AHK4 receptors, as well as in *TCS∷GFP* and *ARR5∷GUS in planta* cytokinin response assays (Fig. S 1a,b and Fig. 1d). This strongly suggested that iP-NBD binds to cytokinin receptors but does not activate them, thus working as a partial cytokinin receptor antagonist. To gain further evidence on selectivity of iP-NBD interactions with cytokinin signalling, we tested its impact on expression of the cytokinin early response gene *ARR5* in plants with different receptor backgrounds. Most pronounced inhibitory effect of iP-NBD on cytokinin’s triggered *ARR5* expression was observed in *ahk2ahk3* double mutant with CRE1/AHK4 receptor functional when compared to either control, *ahk3ahk4* and *ahk2ahk4* seedlings with all, AHK2 and AHK3, receptors active, respectively. This further suggests that iP-NBD might have a higher affinity to CRE1/AHK4 when compared to its homologues (Fig. 1d). Altogether, the above experiments show that iP-NBD exhibits affinity to cytokinin receptors and has potential for specifically tracking their subcellular localization *in planta*.

**Figure 1.**
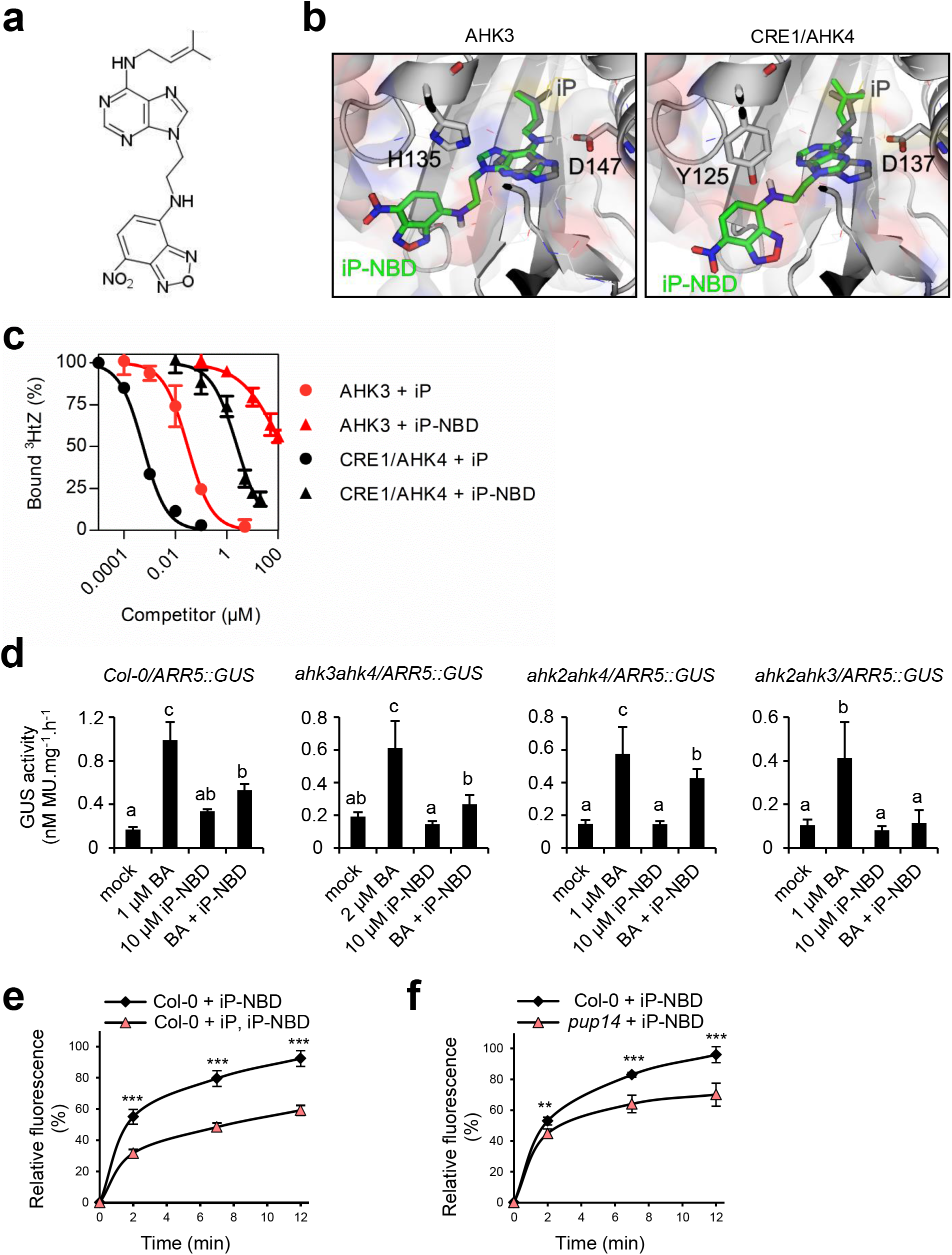
iP-NBD is a cytokinin antagonist interacting with cytokinin perception and transport. **a.** Chemical structure of iP-NBD. **b.** Superposition of docking simulation of iP-NBD (in green) and natural ligand iP (in grey) in AHK3 and CRE1/AHK4 receptor cavity showing the embedded position of the ligands. **c.** Competitive binding assay with *Escherichia coli* expressing AHK3 and CRE1/AHK4. Binding of 3 nM [2-^3^H]*t*Z was assayed together with increasing concentrations of iP-NBD (triangles) and unlabelled iP (circles). The functional inhibition curves for AHK3 and CRE1/AHK4 are presented in red and black, respectively. **d.** Quantitative evaluation of β‐glucuronidase activity in Col-0 and *ahk* double receptor mutants harbouring *ARR5∷GUS* after incubation with cytokinin *N^6^*-benzyladenine (BA), iP-NBD and their combination. Optimal concentration to reach the highest reporter response used was 1 μM, except of *ahk3ahk4* where 2 μM BA was used. The concentration of iP-NBD was 10 μM, mock treatment represents solvent control DMSO (0.1%). The bars represent mean ± s.d., p < 0.01 by ANOVA test, (n=3). MU, 4-methylumbelliferone. **e.** Kinetics of iP-NBD uptake in Arabidopsis LRC cells of wild type (Col-0) pre-treated with iP. **f.** Kinetics of iP-NBD uptake in root meristem epidermal cells of wild type (Col-0) and *pup14*. iP-NBD fluorescence was measured in four time points (0, 2, 7 and 12 min). The bars represent mean ± s.d., *** = p < 0.001, ** = p < 0.01; n ≥ 20 (Student’s t-test) (**e**, **f**).

To reliably monitor iP-NBD distribution *in planta*, we evaluated its biological stability, fluorescence characteristics and saturation kinetics. Degradation of iP-NBD examined by *in vitro* enzymatic reaction with AtCKX2, one of the most active CKX isoforms, revealed that iP-NBD was recognized as the substrate with two-fold slower degradation rate compared to the parental iP molecule (Fig. S1c). To investigate further this potential application constraint, stability of the fluoroprobe was tested *in vivo*. iP-NBD was applied to Arabidopsis cells, and its intracellular processing was followed over a period of 0.5 - 5 h by quantitative liquid chromatography-tandem mass spectrometry (LC-MS/MS) analysis using iP-NBD and *N9*-NBD-labelled adenine (Ade-NBD; the expected product of side-chain cleavage by endogenous CKXs) as molecular standards. Under these conditions, iP-NBD showed high stability within the first 30 minutes (≥ 90% recovery of intact molecule), dropping drastically after 5 hours (Fig. S1d).

In terms of fluorescent characteristics, the emission maximum of the cytokinin fluoroprobe was in the yellow-green part of the spectrum at 528 nm suitable for co-localization with fluorescent markers emitting at red wavelength range (Fig. S1e,g). Quantitative fluorescent microscopy of wild type plants (Col-0) showed that cellular internalization of iP-NBD followed rapid saturation kinetics, reaching a plateau after approximately 12 min (Fig. 1e). Pre-treatment with non-labelled iP and subsequent application of iP-NBD resulted in a significant reduction of intracellular iP-NBD fluorescence (Fig. 1e). This suggested that transport and/or intracellular binding competition between iP-NBD and the natural cytokinin competitor was taking place, further pointing to the cytokinin-like properties of the iP-NBD molecule. Significantly slower progress of iP-NBD accumulation in cells of a *pup14* mutant (lacking the functional cytokinin transporter PUP14) confirmed that specific cytokinin transport partially accounts for the amount iP-NBD detected intracellularly (Fig. 1f). Unlike iP-NBD, Ade-NBD, which lacks the cytokinin-specific side chain, has no affinity to the cytokinin receptors (Fig. S1f) and exhibited a weak diffused apoplastic signal (Fig. S1g).

Affinity of iP-NBD to cytokinin receptors, in particular to CRE1/AHK4, motivated us to monitor subcellular localization of this cytokinin fluoroprobe, aiming to trace potential sites of interaction with the receptor. Two cell types, namely differentiated lateral root cap (LRC) cells and epidermal cells at the root meristematic zone of Arabidopsis root, were selected for in depth analyses. In a line with reported ER-localization of the AHK cytokinin receptors ^2, 3^, iP-NBD co-localized with p24δ5-RFP, an ER-specific marker, in both cell types (Fig. 2a,b, red arrows). Notably, we also detected strong iP-NBD fluorescence signal in distinct spot-like structures, which did not overlap with the ER reporter (Fig. 2a,b; white arrows). Likewise, co-visualization with HDEL-RFP, an ER-specific marker, corroborated both ER and spot-like localization of iP-NBD in cells of LRC (Fig. S2a).

**Figure 2.**
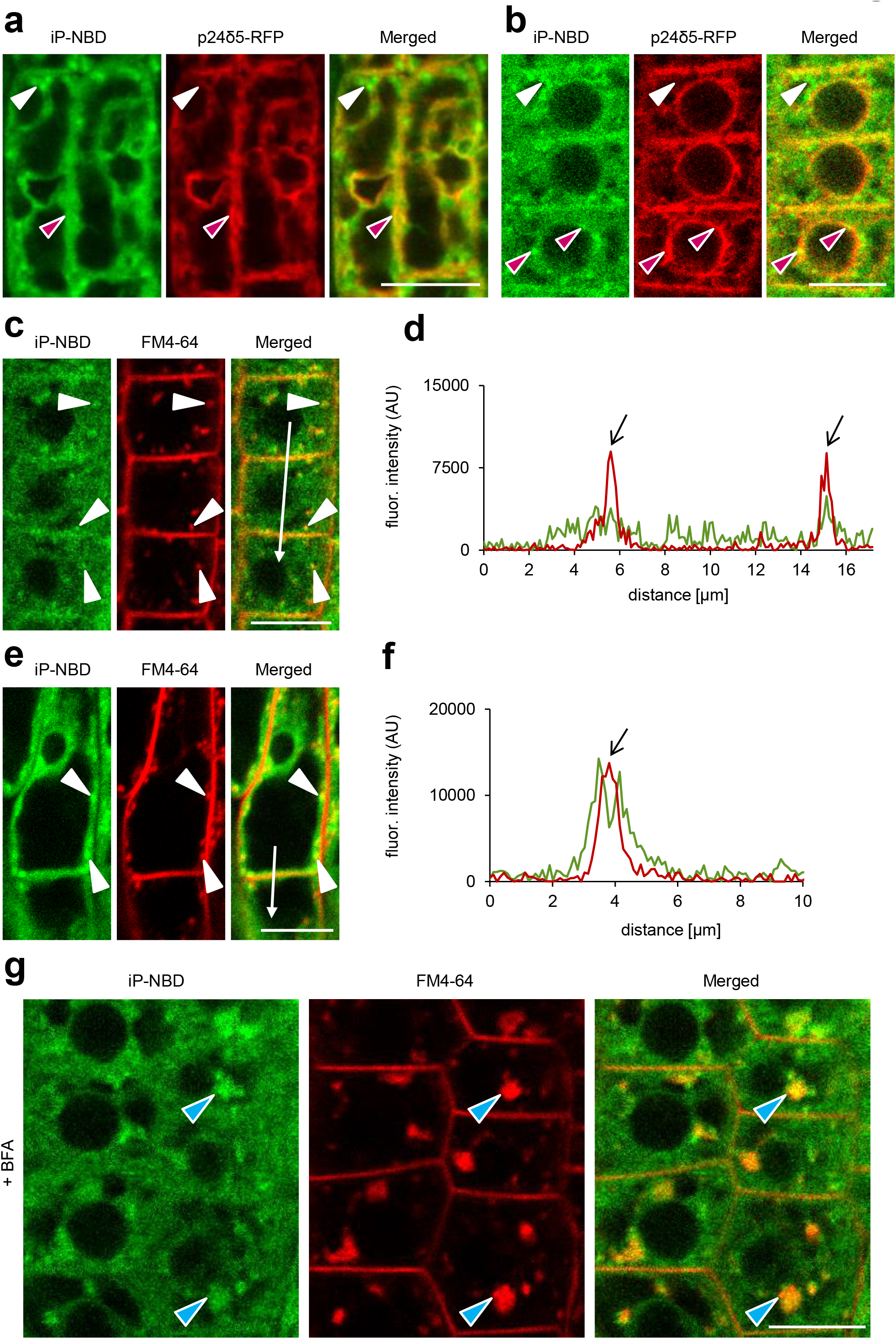
Monitoring of fluorescently labelled cytokinin iP-NBD in cells of Arabidopsis root. **a-b**. Monitoring of fluorescently labelled cytokinin iP-NBD (green) and ER-marker p24δ5-RFP (red) in LRC cells (**a**) and root epidermal cells (**b**). iP-NBD partial co-localization with p24δ5-RFP in ER (red arrowheads) and in non-ER cellular structures (white arrowheads) detected. **c-f.** Monitoring of iP-NBD (green) and FM4-64 (red, membrane selective dye) in root epidermal (**c, d**) and LRC cells (**e, f**). White arrowheads (**c**, **e**) indicate co-localization of iP-NBD and FM4-64 in vesicles. Profiles of fluorescence intensity distribution of both FM4-64 (red line) and iP-NBD (green line) in epidermal (**d**) and LRC (**f**) cells were measured along the white lines (**c**, **d**) starting from upper end (0 μm) towards the arrowhead. Peaks of FM4-64 fluorescence maxima (black arrows) correlate with plasma membrane staining. Peaks of iP-NBD signal partially overlap with FM4-64 maxima and indicate presence of cytokinin fluoroprobe at plasma membrane of epidermal cells (**d**). iP-NBD fluorescence maximum does not overlap with FM4-64 fluorescence peak at plasma membrane in LRC cell (**f**). **g**. Co-localization of iP-NBD and FM4-64 in endosomal compartments (blue arrowheads) formed in root epidermal cells treated with 50 μM in BFA for 1h. Scale bars = 10 μm.

To further explore nature of peripheral and spot-like subcellular structures showing affinity to iP-NBD, we performed co-staining with FM4-64, the membrane selective dye labelling PM and endosomal vesicles of the internalized PM ^13^. In both epidermal cells and LRC, iP-NBD signal was detected intracellularly and partially co-localized with the FM4-64 stained vesicles corresponding to internalized and recycling endosomes (Fig. 2c,e and Fig. S2b). Interestingly, detailed profiles of fluorescence intensity distributions of iP-NBD and FM4-64 revealed their partial co-localization at the PM of epidermal cells, which was not the case for LRC (Fig. 2d compared to 2f). These observations suggested that apart from ER, also PM and subcellular vesicles show high affinity to iP-NBD.

To gain further insights into iP-NBD subcellular localization and to test its affinity to endomembrane structures we analysed impact of brefeldin A (BFA), a fungal toxin, inhibiting ER-Golgi and post-Golgi trafficking to the PM and to vacuoles, thus causing formation of endosomal clusters, so-called BFA compartments ^14^. Strikingly, in root epidermal cells we observed accumulation of iP-NBD signal in clusters corresponding to BFA compartments stained with FM4-64 (Fig. 2g). Co-localization with RabA1e-mCherry, a BFA-sensitive endosome/recycling endosome marker, provided additional supporting evidence that in root epidermal cells iP-NBD exhibits affinity to vesicular endomembrane system where subpopulations of cytokinin receptors may be localized (Fig. S2c). Next, we traced the localization of the cytokinin fluoroprobe using a set of Wave marker lines specific for various subcellular organelles ^15^. Notably, in root epidermal cells we observed a partial co-localization of iP-NBD with a cis-Golgi (GA) marker, SYP32-mCherry (Fig. S2d), an integral GA membrane protein, Got1p homolog-mCherry (Fig. S2e), and with TGN/early endosome marker, VTI12-mCherry (Fig. S2f). Interestingly, iP-NBD did not co-localize with a late endosome marker, RabF2b/W2R-mCherry (Fig. S2g) nor with a vacuolar marker, VAMP711-mCherry (Fig. S2h). In cells of LRC, we observed co-localization with the GA marker, SYP32-mCherry (Fig. S3a) and partial co-localization with the TGN/early endosome marker VTI12 (Fig. S3b). However, no co-localization was detected with endosomal/recycling endosomal RabA1e-mCherry (Fig. S3c), late endosomal RabF2b-mCherry (Fig. S3d) or vacuolar VAMP711-mCherry markers (Fig. S3e).

Overall, monitoring of iP-NBD in LRC and epidermal cells corroborate the ER as an organelle with affinity to cytokinin. However, co-localization of iP-NBD with TGN and early endosomal markers as well as its accumulation in BFA compartments, detected in root epidermal cells, indicate that proteins with affinity to iP-NBD, do not reside exclusively at ER, but may enter endomembrane trafficking system and possibly localize also to the PM.

Previously, ER-localization of Arabidopsis cytokinin receptors has been demonstrated using transiently transformed tobacco (*Nicotiana benthamiana* ^2^,^3^) and Arabidopsis ^3^, and by employing cytokinin-binding assays with fractionated Arabidopsis cells expressing Myc-tagged receptors ^2^. However, so far no experimental support has been provided for their possible entry into the subcellular vesicular trafficking and PM localization. The only cytokinin receptor studied for its localization using stably-transformed Arabidopsis plants was AHK3 ^3^. Unlike this receptor, subcellular localization of CRE1/AHK4 homologue has not been addressed in much detail. Taking into account a higher affinity of iP-NBD to this receptor, we focused on monitoring its subcellular localization using Arabidopsis stable transgenic lines carrying *CRE1/AHK4-GFP* construct driven by a constitutive *35S* promoter. To test functionality of the *CRE1/AHK4-GFP* fusion we performed transient expression assays in Arabidopsis protoplasts. Co-expression of *CRE1/AHK4-GFP* with a cytokinin sensitive reporter *TCS∷LUCIFERASE* (*TCS∷LUC*) resulted in 85 ± 6.9 fold upregulation of the reporter activity by cytokinin when compared to protoplasts co-transformed with either GFP or GUS reporter only resulting in 28 ± 2.4 and 32 ± 1.5 fold increase of LUCIFERASE activity, respectively (Fig. S4a). Moreover, transgenic Arabidopsis seedlings expressing CRE1/AHK4-GFP exhibited phenotypes typical of plants with enhanced activity of cytokinin such as a shorter primary root, slower root growth rate and decreased lateral root density (Fig. S4b-e), indicating that GFP fused to CRE1/AHK4 does not compromise receptor activity.

As previously reported and in line with iP-NBD subcellular localization, CRE1/AHK4-GFP in LRC and epidermal cells of root apical meristem co-localized with ER marker p24δ5-RFP (Fig. 3a,b, red arrows). Intriguingly, in epidermal cells of the root meristematic zone, CRE1/AHK4-GFP signal correlated with the PM reporter PIP1,4-mCherry (Fig. 3c,d). Moreover, in dividing meristematic cells CRE1/AHK4-GFP could also be detected at the expanding cell plate (Fig. 3c,e asterisks) while it co-localized there with established cell plate vesicular marker FM4-64 (Fig. 3f). Importantly, it has been shown that during cytokinesis the cell plate might receive material both from post-Golgi compartments as well as from the PM through sorting and recycling endosomes ^16^. Hence, detection of CRE1/AHK4-GFP at the cell plate provides further supporting evidence that the cytokinin receptor might reside outside of ER, namely on cytokinetic vesicles forming cell plate ^17^. Further evidence confirming localization of CRE1/AHK4-GFP to the PM resulted from the subcellular study using super-resolution structural illumination microscopy (SIM) ^18^. This SIM analysis revealed co-localization of CRE1/AHK4-GFP with FM4-64 labelled PM with average Pearson’s coefficient 0.345 ± 0.113 (Fig. 3g and S5a). Unlike epidermal cells of the root meristematic zone, in LRC cells the CRE1/AHK4-GFP signal resided in the ER and no co-localization with a PM reporter (NPSN12-mCherry) could be detected (Fig. S5b,c). Inhibition of endocytic trafficking and vesicular recycling in meristematic cells by BFA resulted in co-accumulation of CRE1/AHK4-GFP and FM4-64 in the BFA-compartments in line with presence of the receptor in the endomembrane system (Fig. 3h). Wash-out of BFA allowed re-localization of the cytokinin receptor back to the PM indicating that it might cycle between PM and TGN (Fig. S5d). Although occasionally in some cells of LRC co-staining with FM4-64 revealed CRE1/AHK4-GFP in BFA compartments, they were relatively rare and randomly scattered in some LRC cells indicating that CRE1/AHK4-GFP trafficking in differentiated cells of LRC might differ from that observed in epidermal cells of root apical meristem (Fig. S5e). These results indicate that in LRC cells CRE1/AHK4 may reside preferentially at the ER, whereas in epidermal cells of the root apical meristem the cytokinin receptor can enter the endomembrane system and localizes both at the ER and at the PM.

**Figure 3.**
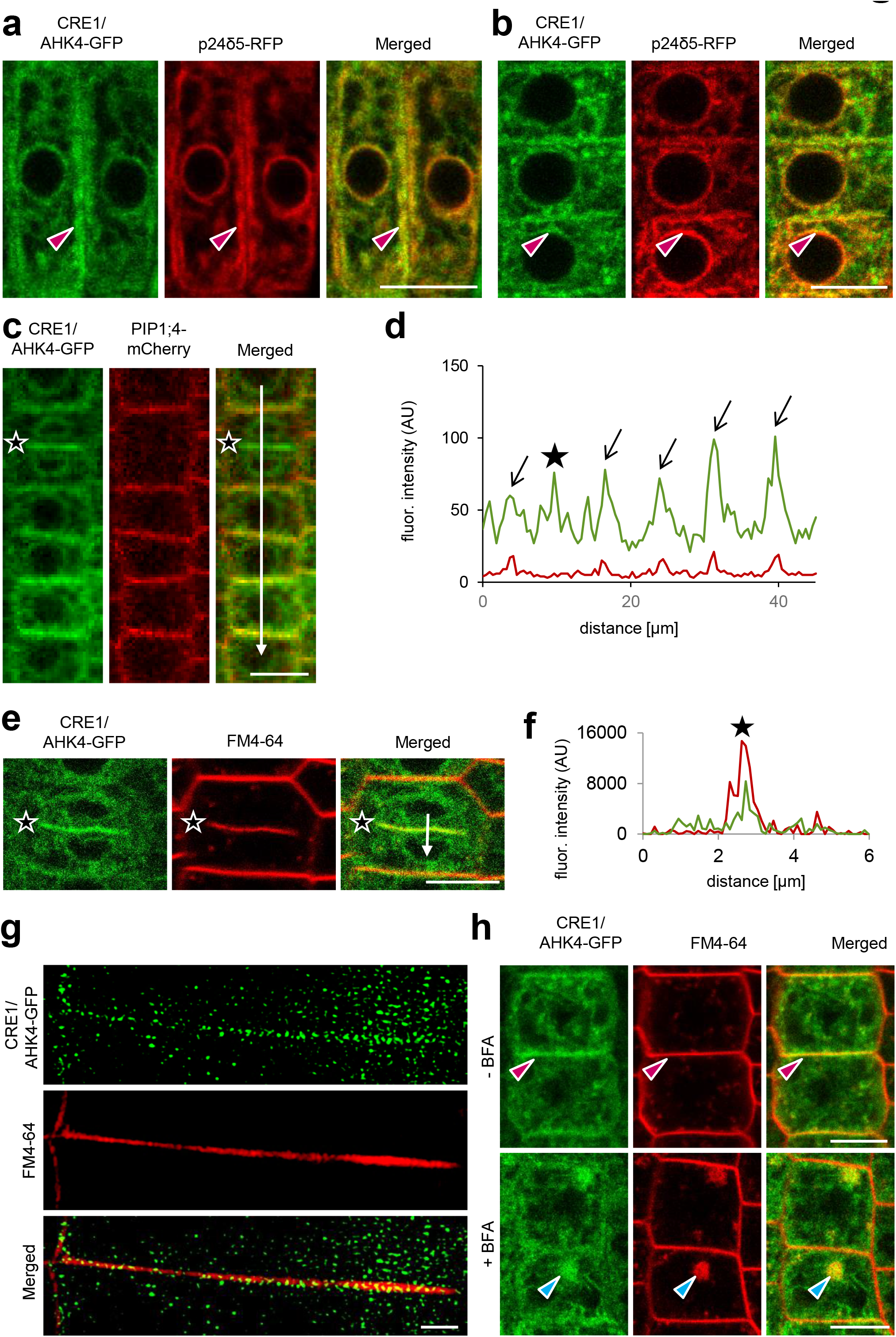
CRE1/AHK4-GFP subcellular localization in cells of Arabidopsis root. **a-b.** Monitoring of CRE1/AHK4-GFP cytokinin receptor (green) and ER-marker p24δ5-RFP (red) in LRC cells (**a**) and root epidermal cells (**b**). Red arrowheads mark areas of co-localization. **c-g**. Co-localization of CRE1/AHK4 with plasma membrane markers PIP1;4-RFP (**c**, **d**) and FM4-64 (**e**-**f**) in the root epidermal cells. Profiles of fluorescence intensity of PM marker (red line) and CRE1/AHK4-GFP (green line) (**d, f**) were measured along the white lines (**c**, **e**) starting from upper end (0 μm) towards the arrowhead. Peaks of PIP1;4-mCherry fluorescence maxima correlate with plasma membrane staining and overlap with CRE1/AHK4-GFP signal maxima (black arrows) (**c**, **d**). CRE1/AHK4-GFP signal detected at the cell plate of dividing cell (black stars on **c**, **e**) and co-localises with FM4-64 marker (**e**, **f**; black stars). (**g**) Super resolution imaging of CRE1/AHK4-GFP subcellular co-localization with FM4-64 labelled PM. (**h)** Co-localization of CRE1/AHK4-GFP and FM4-64 in endosomal compartments (blue arrowheads) formed in root epidermal cells treated with 50 μM in BFA for 1 h. Note CRE1/AHK4-GFP localization at plasma membrane (red arrowheads) prior BFA treatment. Scale bars = 10 μm (a-c, e, h) and 2 μm (g).

Taken together, monitoring of intracellular localization of the fluorescent cytokinin probe iP-NBD with higher affinity to CRE1/AHK4 cytokinin receptor, as well as direct visualization of CRE1/AHK4-GFP leads us to the conclusion that besides ER, cytokinin signal might also be perceived at other cellular compartments including the PM. As suggested by different localization of CRE1/AHK4 receptor in differentiated cells of LRC when compared to epidermal cells of root apical meristem, perception of cytokinin at either ER or PM might be cell- and developmental-context dependent. In particular, the strong expression of the cytokinin sensitive reporter *TCS∷GFP* detected in columella and LRC cells ^19^ suggests that the ER-located cytokinin receptors activate cytokinin signalling cascade in these particular cell types. On the other hand, it remains to be resolved whether there is a specific branch of cytokinin signalling activated by receptors located at the PM of meristematic cells.

## Material and Methods

### Plant material

*Arabidopsis thaliana* (L.) Heynh (Arabidopsis) plants were used. The transgenic lines have been described elsewhere: *cre1-2* ^20^, *pup14* ^5^, p24δ5-RFP ^21^, HDEL-RFP ^22^, Wave lines 2R/RabF2b (LE/PVC), 9R/VAMP711 (Vacuole), 13R/VTI12 (TGN/EE), 18R/Got1p (Golgi), 22R/SYP32 (Golgi), 34R/RabA1e (Endosomal/Recycling endosomal), 131R/NPSN12 (PM) and 138R/PIP1;4 (PM) ^15^, *ARR5∷GUS* ^23^, *ahk3ahk4*/*ARR5∷GUS* ^24^, *ahk2ahk4*/*ARR5∷GUS* ^24^, *ahk2ahk3*/*ARR5∷GUS* ^24^, *TCS∷GFP* ^25^, *35S∷GFP* line was kindly provided by Shutang Tan (IST Austria, Austria). *35S∷CRE1/AHK4-GFP* plants were generated as described below.

### Growth conditions

Surface-sterilized seeds of *Arabidopsis* ecotype Columbia (Col-0) and the other transgenic lines were plated on half-strength (0.5x) Murashige and Skoog (MS) medium (Duchefa) with 1% (w/v) sucrose and 1 % (w/v) agar (pH 5.7). The seeds were stratified for 2-3 days at 4 °C in darkness. Seedlings were grown on vertically oriented plates in growth chambers at 21 °C under long day conditions (16 h light and 8 h darkness) using white light (W), which was provided by blue and red LEDs (70-100 μmol m^−2^s^−1^ of photosynthetically active radiation), if not stated otherwise.

### Pharmacological and hormonal treatments for microscopy imaging

Seedlings 4-5 days old were transferred onto solid 0.5x MS medium with or without the indicated chemicals. The drugs and hormones used were: *N^6^*-benzyladenine (BA) in different concentrations (0.1 μM, 0.5 μM, 1 μM and 2 μM), *trans*-zeatin (*t*Z 0.1 and 1 μM), *N^6^*-isopentenyladenin (iP, 5 μM). Mock treatments were performed with equal amounts of solvent (DMSO). Treatments with 5 μM iP-NBD and 5 μM Ade-NBD were performed in liquid 0.5x MS medium and imaging was carried out within 30 min time frame. For co-localization of the cytokinin fluoroprobe with PM marker, seedlings were pre-treated with 2 μM FM4-64 for 5 min and transferred into 5 μM iP-NBD supplemented 0.5x MS medium, followed by imaging within 30 minutes time frame. To explore affinity of iP-NBD to BFA endosomal compartments, seedlings were incubated in 50 μM BFA for 1 h and afterwards transferred into iP-NBD supplemented medium and imaged. Localization of CRE1/AHK4-GFP in BFA endosomal compartments was examined in 4-5 days old seedlings incubated in 50 μM BFA for 1 h. For BFA washout experiments, seedlings were placed in a fresh BFA-free 0.5x MS medium, which was replaced every 10 min for at least 1 h.

### Recombinant DNA Techniques

The coding region of the cytokinin receptor CRE1/AHK4 (*At2g01830*) was amplified without the stop codon by PCR using a gDNA from *Arabidopsis thaliana* Col-0 as a template and cloned into the Gateway vector *pENTR_2B* dual selection (primers: AHK4_Fw_SalI_KOZAK *CGCGTCGACccaccATGAGAAGAGATTTTGTGTATAATAATAATGC* and AHK4_R_NotI *CCTAATCCTATCTCACCTTCGTCGtcGCGGCCGCAAAAGGAAAA*). To construct C-terminal fusion of *CRE1/AHK4* with *GFP*, *CRE1/AHK4* was shuttled into the destination vector *pGWB5* containing *35S* promoter to create *35S∷AHK4-GFP* construct. For the transient Luciferase assay in Arabidopsis protoplasts, *AHK4-GFP* fusion construct was re-cloned into *p2GW7,0* vector. *AHK4-GFP* region was amplified by PCR using *pGWB5_AHK4-GFP* as a template (primers: *35S_FW CCACTATCCTTCGCAAGAC*CCTTC and AHK4_5A_NheI_RE *TATTCCAATgctagcTTACTTGTACAGCTCGTCCATGC*) and ligated into the Gateway *pENTR_2B* dual selection entry vector. *CRE1/AHK4-GFP* was shuttled into the destination vector *p2GW7,0*.

### Plant transformation

Transgenic Arabidopsis plants were generated by the floral dip method using *Agrobacterium tumefaciens* strain GV3101 ^26^. Transformed seedlings were selected on medium supplemented with 30 μg mL^−1^ hygromycin.

### Homology modelling and molecular docking

CRE/AHK structures were modelled based on CRE1/AHK4-iP crystal structure (PDBID: 3T4L) ^9^ using Modeller 9.10 ^27^. The geometry of iP-NBD was modelled with Marvin (http://www.chemaxon.com), and then the compounds were prepared for docking in the AutoDockTools program suite ^28^. The Autodock Vina program ^29^ was used for docking iP-NBD ligand into the set of CRE/AHK structures obtained from homology modelling. A 15 Å box centred at the original ligand binding position was used. The exhaustiveness parameter was set to 20.

### Competitive binding assay with CRE1⁄AHK4- and AHK3-expressing *E. coli* strains

Receptor direct binding assays were conducted using the *E. coli* strain KMI001 harboring either the plasmid *pIN-III-AHK4* or *pSTV28-AHK3*, which express the Arabidopsis histidine kinases CRE1/AHK4 or AHK3 ^30, 31^. Bacterial strains were kindly provided by Dr. T. Mizuno (Nagoya, Japan). The assays were performed as described in ^12^ with slight modifications presented in ^32^. 3 nM [2-^3^H]*t*Z was used as the radiolabeled natural ligand. The functional inhibition curves were used to estimate the IC_50_ values and the K_i_ values were calculated according to Cheng and Prusoff equation ^33^. [2-^3^H]*t*Z was provided by Dr. Zahajská from the Isotope Laboratory, Institute of Experimental Botany ASCR.

### *ARR5∷GUS* reporter gene assay

Arabidopsis *ARR5∷GUS* transgenic seeds were surface-sterilized and plated on 0.5x MS medium with 0.1% (w/v) sucrose and 0.05% (w/v) MES– KOH (pH 5.7) in 24-well plates. The seeds were stratified for 3 days at 4 °C in darkness and then grown under long-day conditions (16 h light/8 h dark) at 22 °C in a growth chamber. To the wells containing 3-day-old seedlings, BA and/or tested compound or DMSO (solvent control, final concentration 0.1%) was added and the seedlings were grown for an additional 16 h, as published previously ^32^. Quantitative determination of GUS activity was performed by measuring fluorescence using Synergy H4 Multi-Mode Microplate Reader (BioTek, USA) at excitation and emission wavelengths of 365 and 450 nm, respectively. Protein content determination was done by bicinchonin acid method and GUS specific activity was expressed as nmol 4-methylumbelliferone (MU)/h/mg protein.

### Measurements of iP-NBD cell transport kinetics

Seedlings of 4-day-old Arabidopsis Col-0 were pre-treated for 20 min with 5 μM iP or DMSO and transferred into MS media containing 5 μM iP-NBD and 5 μM iP/DMSO and instantly imaged. To examine PUP14 dependent iP-NBD transport kinetics, 4-day-old seedlings of *pup14* and Col-0 were treated with 5 μM iP-NBD and immediately imaged. For both experiments imaging was performed in the same area of the root for 12 min every 2, 7 and 12 min to minimize photobleaching. iP-NBD fluorescence was measured with ImageJ in the LRC cells (iP pre-treatment experiment) and in the root epidermal cells (*pup14* experiment) from 4-7 cells originating from 4-5 roots.

### Imaging

For confocal microscopy imaging, a vertical-stage laser scanning confocal Zeiss 700 (LSM 700) and Zeiss 800 (LSM 800), described in ^34^, with a 20×/0.8 Plan-Apochromat M27 objective and a LSM 800 inverted confocal scanning microscope Zeiss, with a 40× Plan-Apochromat water immersion objective, were used. Samples were imaged with excitation lasers 488 nm for GFP (emission spectrum 490-560 nm) and NBD (emission spectrum 529-570 nm) and 555/561 nm (inverted/vertical) for RFP (emission spectrum 583-700 nm), FM4-64 (emission spectrum 650-730 nm) and mCherry (emission spectrum 570-700 nm).

For super-resolution SIM microscopy, an Axioimager Z.1 with Elyra PS.1 system coupled with a PCO.Edge 5.5 sCMOS camera was used. Samples were excited with the 488nm and 561nm laser lines. Oil immersion objective (63×/1.40) and standard settings (the grating pattern with 5 rotations and 5 standard phase shifts per angular position) were used for image acquisition. Image reconstruction was done according to the published protocol ^35^. For image post-processing, profile measurements and co-localization analysis, the Zeiss Zen 2011, ImageJ (National Institute of Health, http://rsb.info.nih.gov/ij), Photoshop 6.0/CS, GraphPad Prism 8 and Microsoft PowerPoint programs were used. For SIM co-localization experiments 30 PM regions originating from root cells of 5 seedling plants were used.

### Statistics

The statistical significance was evaluated with the Student’s t-test and ANOVA.

## Author contributions

LS, OP, EB, PG and MS conceived the project; KK and JCM performed most of the confocal and biochemical experiments; OŠ and JŠ participated in plasma membrane localization studies and helped with evaluation and interpretation of subcellular localization data; LP, KD and VM designed and chemically synthesized cytokinin fluorescent probe; JN performed cytokinin in vitro assays; ON conducted the purification and quantification of cytokinins; KB performed *in silico* docking experiments; DZ analysed and interpreted the data; LS, EB and OP designed experiments, analysed and interpreted the data; KK, JCM, OŠ and LS made the Figures; EB, OP and LS wrote the paper.

## Acknowledgements

This paper is dedicated to deceased P. Galuszka for his support and contribution to the project. This research was supported by the Scientific Service Units (SSU) of IST-Austria through resources provided by the Bioimaging Facility (BIF), the Life Science Facility (LSF) and by Centre of the Region Haná (CRH), Palacký University. We thank Hana Semerádová and Lucia Hlusková for technical assistance, and Fernando Aniento, Rashed Abualia and Andrej Hurný for sharing material. The work was supported from ERDF project "Plants as a tool for sustainable global development" (No. CZ.02.1.01/0.0/0.0/16_019/0000827), from Czech Science Foundation via projects 16-04184S (OP, KK and KD), 18-23972Y (DZ), 17-21122S (KB), IGA_PrF_2017_016 (KK), and EMBO Long-Term Fellowship, ALTF number 710-2016 (JCM). The authors declare that they have no conflicts of interest.

## Supplementary Information

**Figure S1.**
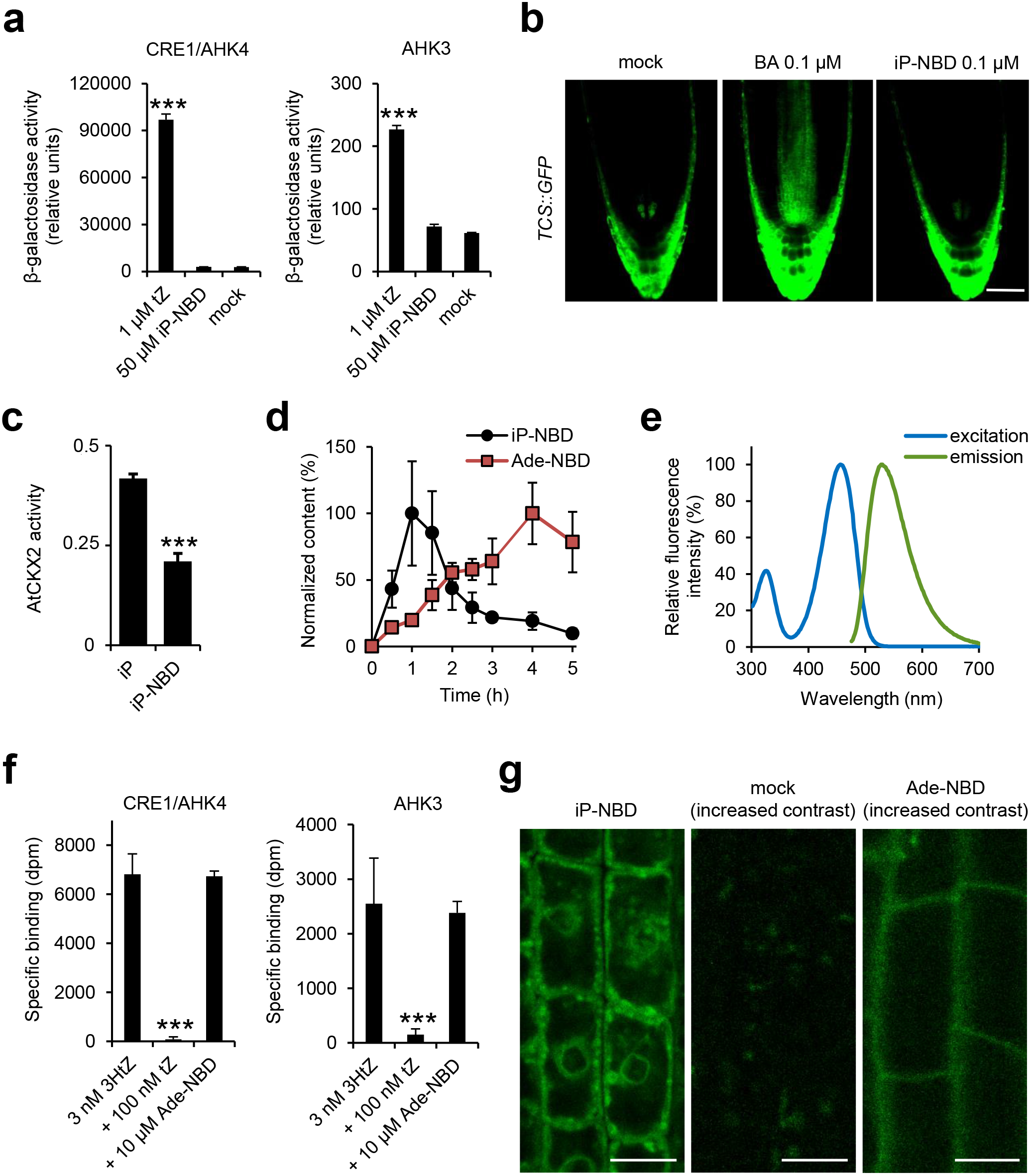
Biological characterization of iP-NBD. **a.** Comparison of cytokinin response in *E. coli* (*ΔrcsC, cps∷lacZ*) receptor activation assay with recombinant CRE1/AHK4 and AHK3 receptors, triggered by 1 μM *t*Z (positive control) and 50 μM iP-NBD (mean ± s.d., *** = p < 0.001 by Student’s t-test, n = 3). **b.** Comparison of the effect of 0.1 μM *N^6^*-benzyladenine (BA; positive control) and 0.1 μM iP-NBD on induction of the cytokinin reporter *TCS∷GFP* in Arabidopsis roots after 15 h treatment. Scale bar = 50 μm. **c.** *In vitro* enzymatic activity of AtCKX2 estimated using 150 nM iP and iP-NBD as substrates, respectively (mean ± s.d., *** = p < 0.001 by Student’s t-test, n = 3). **d.** Stability of iP-NBD *in vivo*. iP-NBD was applied to *Arabidopsis* (L*er*) cells suspension and in the timeframe of 0.5-5 h its intracellular processing was followed by quantitative LC-MS/MS analysis using iP-NBD and Ade-NBD (the expected product of side-chain cleavage by endogenous CKXs) as molecular standards (as described in the methods). The values presented in the graph are normalized to respective highest content of both compounds analysed. The highest contents of intracellular iP-NBD and Ade-NBD were 7920 ± 3150 pmol/g and 19633 ± 4903 pmol/g, respectively (FW, mean ± s.d., n = 4). **e.** Fluoroprobe absorption-emission spectral diagram measured with 100 μM iP-NBD dissolved in 100% ethanol. Absorption (excitation) and emission spectra are reaching their maximal fluorescence intensities in 456 nm (ex) and 528 nm (em). **f.** Competitive binding assay with *Escherichia coli* expressing AHK3 and CRE1/AHK4 with Ade-NBD. Binding of 3 nM [2-^3^H]*t*Z was assayed together with high excess concentration of Ade-NBD (10 μM), and unlabelled *t*Z (100 nM) as a positive control (mean ± s.d., *** = p < 0.001 by Student’s t-test, n = 3). **g.** Differential localization of iP-NBD and Ade-NBD in Arabidopsis LRC cells. Roots were treated for 10 min with iP-NBD or Ade-NBD (5 μM). Roots without any treatment (mock) were used as a control. Scale bar = 10 μm.

**Figure S2.**
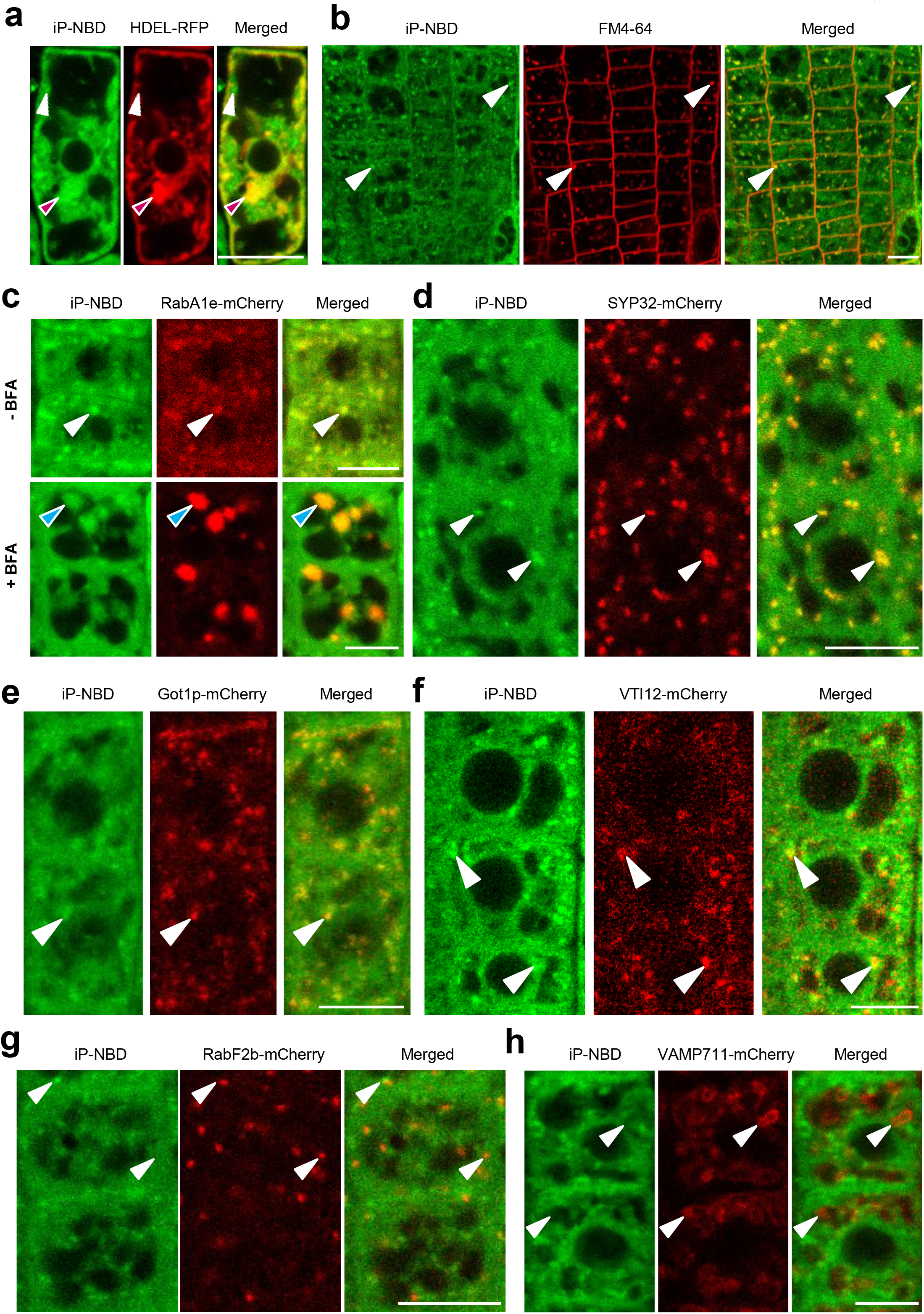
Monitoring of iP-NBD subcellular localization in Arabidopsis cells. **a.** Monitoring of iP-NBD (green) and ER-marker HDEL-RFP (red) in LRC cells. iP-NBD partial co-localization with HDEL-RFP in ER (red arrowheads) and in non-ER cellular structures (white arrowheads) detected. **b.** Co-staining of root epidermal cells with iP-NBD (green) and FM4-64 (red). White arrowheads indicate co-localization of iP-NBD and FM4-64 in vesicles. **c.** Co-localization of iP-NBD (green) and RabA1e-mCherry (red) endosome/recycling endosome marker. Upper panel: co-localization of iP-NBD with RabA1e in vesicles (white arrowheads) before BFA treatment. Lower panel: accumulation of iP-NBD and RabA1e in the endosomal compartments (blue arrowheads) formed in root epidermal cells treated with 50 μM BFA for 1 h. **d-f**. Partial co-localization of iP-NBD (green) with a cis-GA marker SYP32-mCherry (**d**), an integral GA membrane protein Got1p-mCherry (**e**) and TGN/early endosome marker VTI12-mCherry (**f**) in root epidermal cells. White arrowheads indicate overlapping signals. **g, h**. Non-overlapping signals of iP-NBD (green) and late endosome marker RabF2b-mCherry (red, **g**) or vacuolar marker VAMP711-mCherry (red, **h**) in root epidermal cells. White arrowheads indicate RabF2-mCherry stained endosomes (**g**) and VAMP711-mCherry vacuolar compartments (**h**). Scale bars = 10 μm.

**Figure S3.**
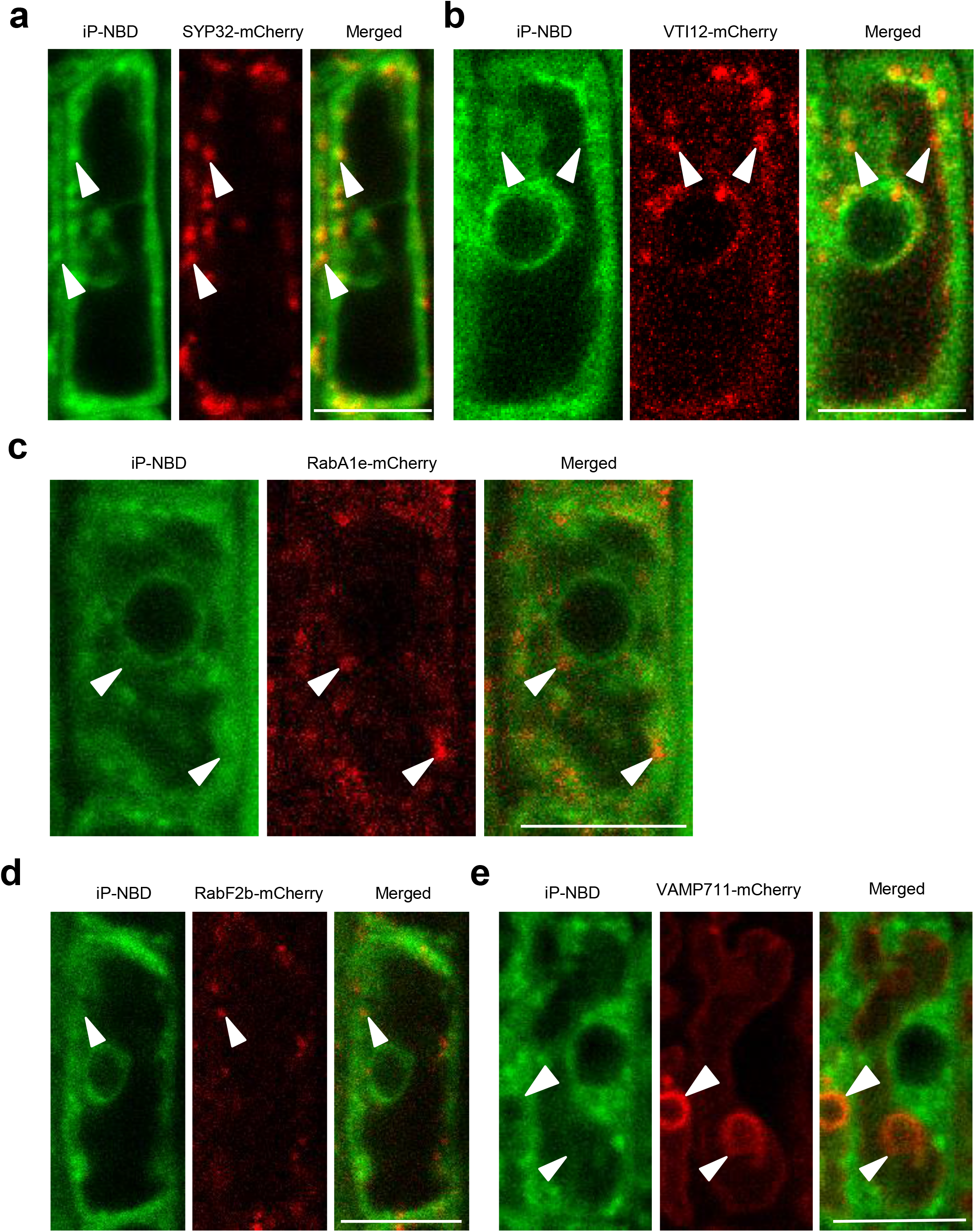
Monitoring of iP-NBD subcellular localization in Arabidopsis lateral root cap cells. **a.** Co-localization of iP-NBD (green) with a cis-GA marker SYP32-mCherry (red) in LRC cell. White arrowheads indicate overlapping signals. **b-e**. Non-overlapping signals of iP-NBD (green) and TGN/early endosome marker VTI12-mCherry (**b**), endosome/recycling endosome marker expressing RabA1e-mCherry (**c**), late endosome marker RabF2b-mCherry (**d**) or vacuolar marker VAMP711-mCherry (**e**, red) in LRC cells. White arrowheads indicate subcellular compartments visualised by specific markers. Scale bars = 10 μm.

**Figure S4.**
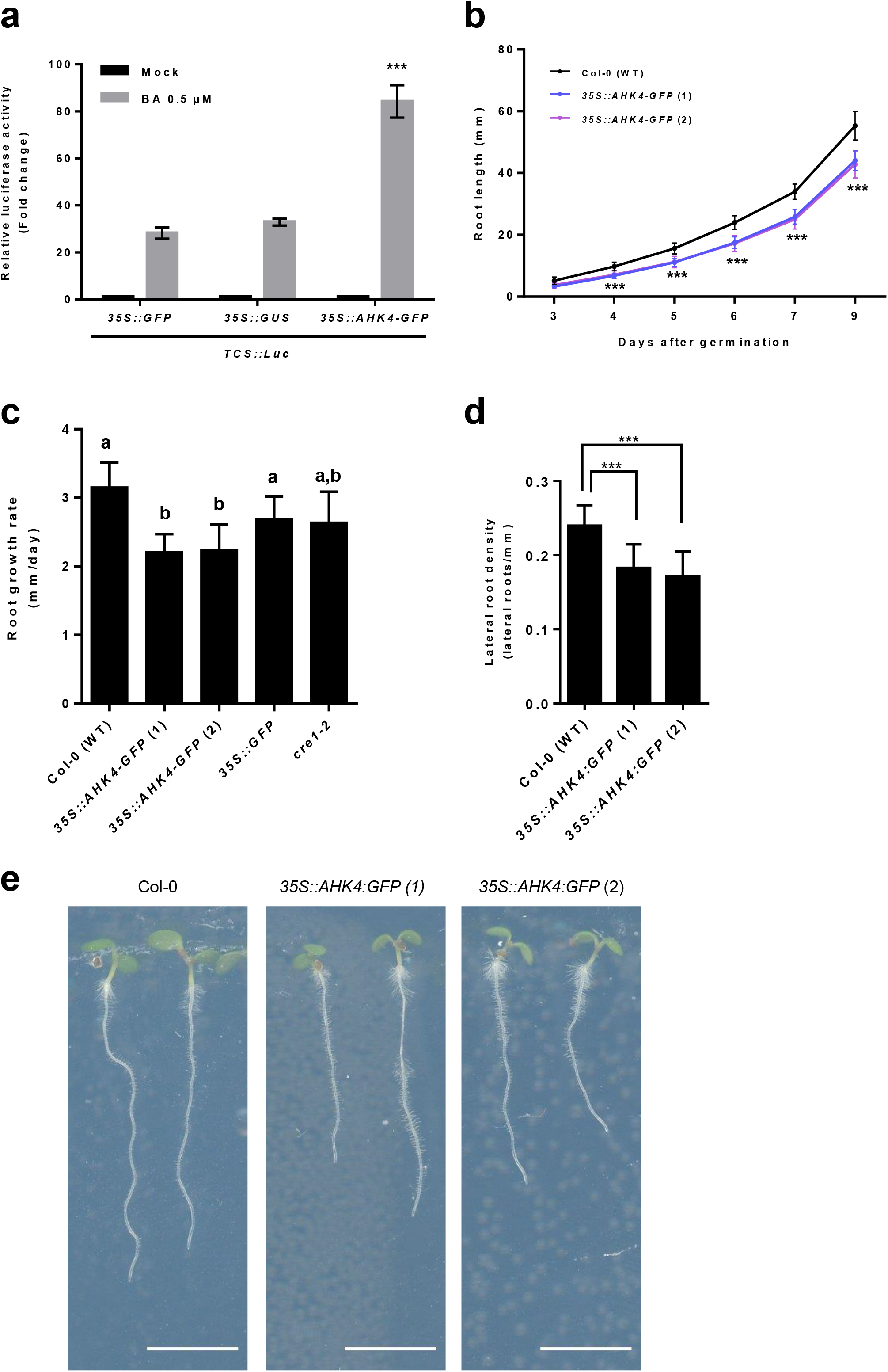
Analysis of CRE1/AHK4-GFP functionality *in vivo*. **a**. *TCS∷LUCIFERASE* cytokinin reporter activity in *Arabidopsis* protoplasts co-transformed with CRE1/AHK4-GFP reporter is significantly upregulated in response to cytokinin (0.5 μM BA) when compared to protoplasts co-transformed with either GFP or GUS reporter only (mean + s.d; *** p < 0.001 Student’s t-test indicates significant difference when compared to protoplasts transformed with *35S∷GUS*, n = 4). **b.** Root length of 3- to 9-day-old seedlings of control Col-0 (black) and two independent *35S∷AHK4-GFP* lines (mean ± s.d., *** = p < 0.001 by Student’s t-test, n ≥15). **c.** Average root growth rate (mm/day) of control Col-0, two independent *35S∷AHK4-GFP* lines *35S∷GFP* and *cre1-2* seedlings during 5 days (mean ± s.d.; p < 0.01 by ANOVA test, n ≥ 15). **d.** Lateral root density (number of lateral roots/root length) was evaluated in 9-day-old seedlings of control Col-0 and two independent *35S∷AHK4-GFP* lines (mean ± s.d.; *** = p < 0.001 by Student’s t-test, n ≥15). **e.** Representative pictures of 5-day-old seedlings of control Col-0 and two independent *35S∷AHK4-GFP* lines. Scale bar = 5 mm.

**Figure S5.**
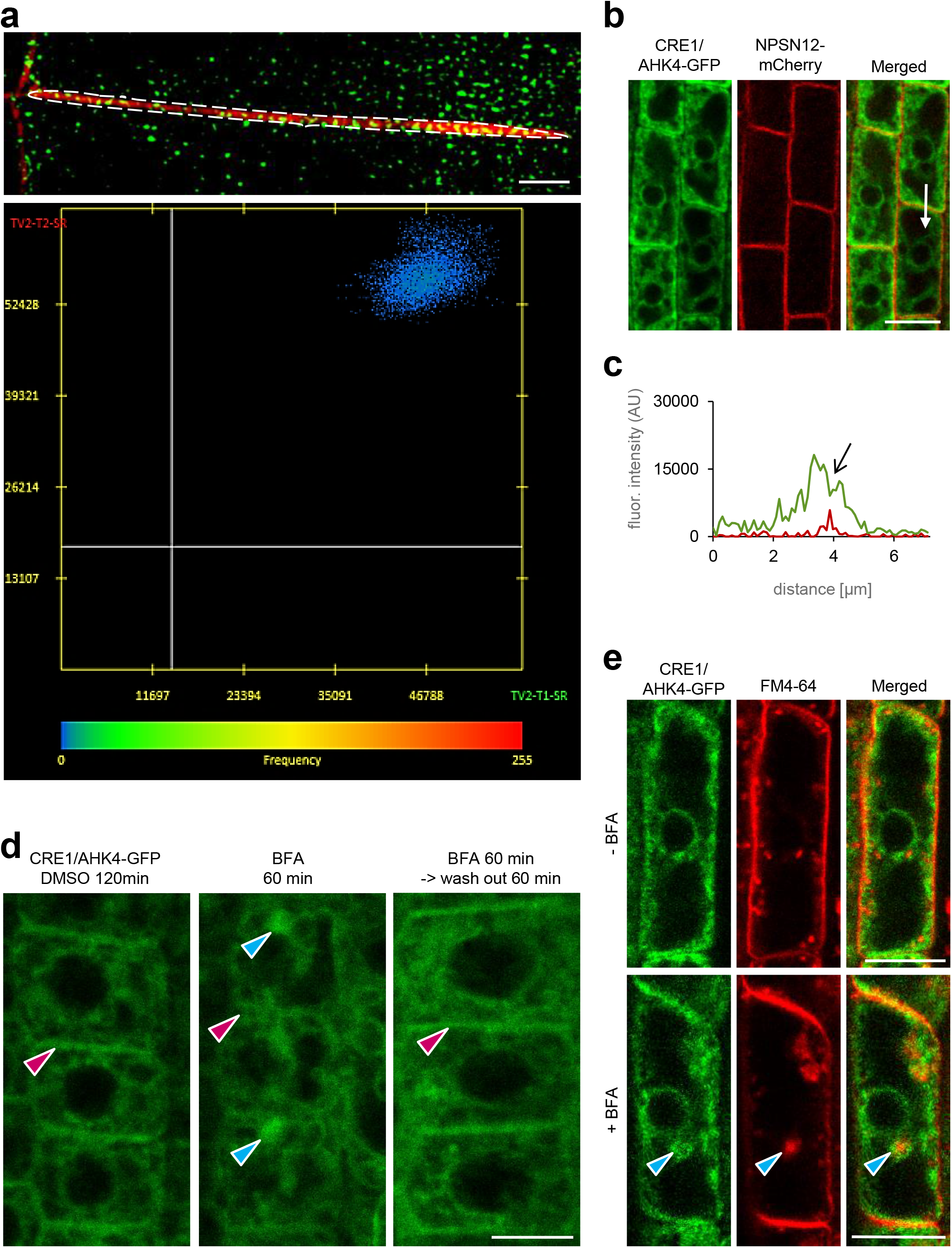
CRE1/AHK4-GFP localization in Arabidopsis root cells. **a.** Co-localization analysis of CRE1/AHK4 with FM4-64 labelled PM in epidermal cells of the root meristem. Dotted line in upper image corresponded to PM co-localization area. Bottom image shows scatter plot of marked PM area in upper image. **b, c.** Monitoring of CRE1/AHK4-GFP cytokinin receptor (green) and NPSN12-RFP plasma membrane reporter in LRC cells (**b**). Profiles of fluorescence intensity of plasma membrane marker (red line) and CRE1/AHK4-GFP (green line) were measured along the white line (**b**) starting from upper end (0 μm) towards the arrowhead. Peak of NPSN12-mCherry fluorescence maxima (black arrow) correlates with plasma membrane staining. CRE1/AHK4-GFP fluorescence maximum does not overlap with NPSN12-mCherry fluorescence peak at plasma membrane (**c**). **d.** Re-location of CRE1/AHK4-GFP cytokinin receptor from the endosomal compartments to the plasma membrane after BFA wash-out in the root epidermal cells. Blue arrowheads indicate CRE1/AHK4 in the BFA-bodies. Note the attenuated plasma membrane signal and its re-localization back after 60 min wash-out of BFA (red arrowheads). **e.** Localization CRE1/AHK4-GFP and FM4-64 in LRC cells treated for 1 h with 50 μM BFA. Blue arrowheads indicate the BFA endosomal compartments stained with FM4-64. Scale bars = 10 μm (b, d, e) and 2 μm (a).

**Figure S6.**
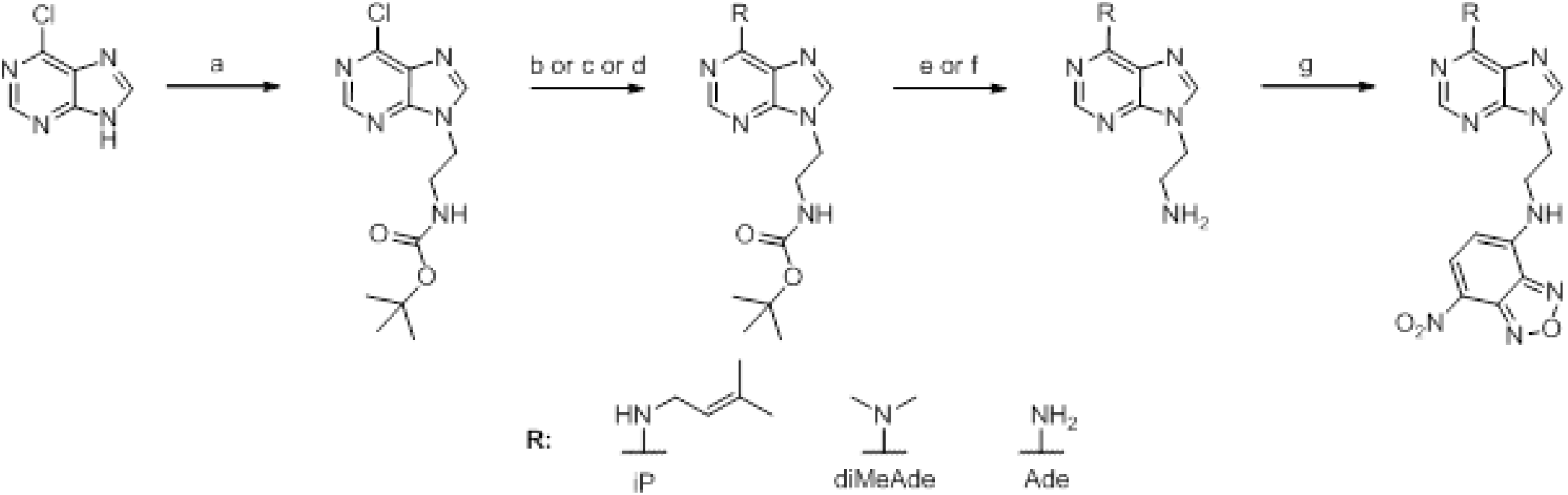
Reaction scheme for preparation of fluorescently labelled probes for confocal microscopy studies. a) *N*-Boc-ethanolamine, PPh_3_, DIAD, THF, rt, 2 h, (68%); b) 3-methylbut-2-enylamine.HCl, Et_3_N, *n*PrOH, reflux, 4 h, (93%); c) Me2NH.HCl, Et_3_N, *n*PrOH, reflux, 4 h, (97%); d) ammonium hydroxide, EtOH, 95 °C, overnight, (76%); e) Dowex 50W X8, DCM, reflux followed by 4 M methanolic ammonia, rt, overnight, (iP - 90%, diMeAde - 87%); f) TFA, DCM, rt, overnight, (Ade.TFA salt - 95%); g) 4-chloro-7-nitro-1,2,3-benzoxadiazole, NaHCO_3_, MeOH, 50 °C, 1 h followed by rt overnight (iP - 52%, diMeAde - 75%, Ade - 68%).

## Supplementary Methods

### Receptor activation assay

The receptor activation assays were conducted using the *E. coli* strain KMI001 harbouring either the plasmid *pIN-III-AHK4* or *pSTV28-AHK3*, which express the Arabidopsis histidine kinases CRE1/AHK4 or AHK3 ^30, 31^. Bacterial strains were kindly provided by Dr. T. Mizuno (Nagoya, Japan). The assays were performed following protocol described in ^36^.

### *TCS∷GFP* reporter expression in planta

5-day-old *TCS∷GFP* seedlings were transferred on non- or with either cytokinin (0.1 μM BA) or 0.1 μM iP-NBD supplemented Murashige and Skoog media for 15 h. *TCS∷GFP* expression (green) in root tip was monitored using confocal microscope.

### AtCKX2 activity measurement

The recombinant enzyme AtCKX2 and the enzymatic activity determination was performed as described in ^37^, using 150 nM iP or iP-NBD as substrates.

### LC-MS/MS method

To study iP-NBD stability *in vivo* 5 μM iP-NBD was applied to *Arabidopsis* (L*er*) cell suspension and in the timeframe of 0.5 - 5 h samples were taken and deep-frozen for subsequent quantitative LC-MS/MS analysis of iP-NBD and Ade-NBD (the expected product of side-chain cleavage by endogenous CKXs) contents. The samples (around 50 mg fresh weight) were extracted in 1.0 mL of modified Bieleski buffer (60% MeOH, 10% HCOOH and 30% H_2_O; ^38^ together with 0.01 pmol of N9-NBD-labelled dimethyladenine (diMeAde-NBD) added as internal standard to validate LC-MS/MS determination. The extracts were purified using the Oasis MCX column (30 mg/1 mL, Waters) and targeted analytes were eluted using 0.35 M NH_4_OH in 60% (v/v) MeOH solution^39^. The purified samples were eluted using a reversed-phase column (Acquity UPLC® BEHC18, 1.7 μm, 2.1 × 150 mm, Waters) with a 26 min gradient comprised of methanol (A) and 15 mM ammonium formate pH 4.0 (B) at a flow rate of 0.25 mL/min and column temperature of 40 °C ^40^. The binary linear gradient of 0-7 min 5:95 A:B, 16 min 20:80 A:B, 23 min 50:50 and 26 min 100:0 was used, after which the column was washed with 100% methanol for 1 min and re-equilibrated to initial conditions for 3 min. The effluent was introduced into the MS system with the following optimal settings: source/desolvation temperature 150/600 °C, cone/desolvation gas flow 150/1000 l h^−1^, capillary/cone voltage 3500/25 V, collision energy 20 eV and collision gas flow (argon) 0.15 mL min^−1^. Quantification and confirmation of the NBD-labelled compounds were obtained by the multiple reaction monitoring mode using the following mass transitions: 410>342, 342>136 and 370>164 for iP-NBD, Ade-NBD and diMeAde-NBD, respectively. All chromatograms were analyzed with MassLynx software (version 4.1; Waters Corporation) and the compounds were quantified according to the internal standard added.

### Estimation of iP-NBD fluorescence properties

Fluorescence spectrum reaching the maximal fluorescence intensities in 456 nm (ex) and 528 nm (em) was obtained with 100 μM iP-NBD dissolved in 100% ethanol by relative fluorescence intensity scanning in the range of 300-700 nm using Synergy H4 Multi-Mode Microplate Reader (BioTek, USA).

### Luciferase transient expression assay

Protoplasts were isolated from root suspension culture and the *Luciferase* transient expression assays performed as previously described ^41^ with slight modifications. Protoplasts were co-transfected with 3 μg of a reporter plasmid expressing *Firefly* luciferase, 2 μg of normalization plasmid containing the *Renilla* luciferase and 10 μg of *p2GW7,0* plasmid carrying either cytokinin receptor (*35S∷AHK4-GFP*) or reporter only (*35S∷GUS* or *35S∷GFP*) constructs. Transfected protoplasts were incubated for 16 h with either 0.5 μM BA or in mock solution. The mean value was calculated from four measurements and experiment was repeated three times.

### Root growth analysis

Root growth and lateral root density were measured on seedlings grown vertically on Murashige and Skoog medium. Images were taken with a vertically positioned scanner, EPSON perfection v800 Photo. Root growth rate was evaluated using 5-day-old seedlings (n ≥ 16). Lateral root density quantification was performed using 9 days-old seedlings (n ≥ 16). The statistical significance was evaluated with the Student’s t-test and ANOVA.

### Synthesis of fluorescently-labelled compounds

The fluorescently labelled compounds used in the studies were prepared according to the reaction scheme (Fig. S6). Two carbon linker terminated with amino group suitable for consecutive NBD fluorophore attachment was coupled to the purine *N*9 position by the reaction of 6-chloropurine with *N*-Boc-ethanolamine under Mitsunobu conditions ^7^. Later, nucleophilic substitution of C6 chlorine with appropriate amines in boiling alcohols followed by Boc protective group cleavage provided purine intermediates for fluorescent labelling. 4-chloro-7-nitro-1,2,3-benzoxadiazol was used as NBD donor and was linked to the purine primary amino group in MeOH under basic conditions. Physico-chemical characterization of the synthesized compounds was done according to ^7^.

### *tert*-butyl [2-(6-amino-9*H*-purin-9-yl)ethyl]carbamate

*tert*-butyl [2-(6-amino-9*H*-purin-9-yl)ethyl]carbamate (0.5 g, 1.68 mmol) was heated in closed vessel with ammonium hydroxide (5 mL) and EtOH (5 mL) at 95 °C overnight. After evaporation of solvents the residue was purified by silica column chromatography using CHCl_3_/MeOH (9:1, v/v) as a mobile phase. White solid, yield 76%. HPLC purity 99.9, ESI^+^-MS 279 (100, [M+H]^+^), ^1^H-NMR (500 MHz, DMSO-*d*_6_) δ (ppm): 1.30 (s, 9H, Boc (C<u>H</u>_3_)_3_), 3.32 (q, *J* = 5.9 Hz, 2H, CH_2_C<u>H</u>_2_NHBoc), 4.16 (t, *J* = 6.0 Hz, 2H, C<u>H</u>_2_CH_2_NHBoc), 6.96 (t, *J* = 5.7 Hz, 1H, CH_2_CH_2_N<u>H</u>Boc), 7.15 (s, 2H, Ade NH_2_), 7.99 (s, 1H, pur H8), 8.11 (s, 1H, pur H2). ^13^C-NMR (125 MHz, DMSO-*d*_6_) δ (ppm): 28.1 (Boc (C<u>H</u>_3_)_3_C), 39.2 (HMQC based, CH_2_<u>C</u>H_2_NHBoc), 42.7 (<u>C</u>H_2_CH_2_NHBoc), 77.8 (Boc (CH_3_)_3_<u>C</u>), 118.7 (pur C5), 140.8 (pur C8), 149.6 (pur C4), 152.2 (pur C2), 155.5 (Boc <u>C</u>O), 155.9 (pur C6).

### 9-(2-aminoethyl)adenine trifluoroacetate

*tert*-butyl [2-(6-amino-9*H*-purin-9-yl)ethyl]carbamate (0.35 g, 1.26 mmol) was added to a mixture of DCM (10 mL) and TFA (0.5 mL, 6.54 mmol) and stirred at room temperature overnight. Reaction mixture was evaporated under reduced pressure and the residue was treated with Et_2_O to obtain white solid. Yield 95%, HPLC purity 99.9, ESI^+^-MS 179 (100, [M+H]^+^), ^1^H-NMR (500 MHz, DMSO-*d*_6_) δ (ppm): 3.36 (s, 2H, CH_2_C<u>H</u>_2_NH_2_), 4.47 (t, J = 5.7 Hz, 2H, C<u>H</u>_2_CH_2_NH_2_), 8.13 (s, 3H, CH_2_CH_2_N<u>H</u>_3_^+^), 8.34 (s, 1H, pur H8), 8.43 (s, 1H, pur H2), 8.72 (s, 2H, Ade NH_2_). ^13^C-NMR (125 MHz, DMSO-*d*_6_) δ (ppm): 38.3 (CH_2_<u>C</u>H_2_NH_2_), 41.4 (<u>C</u>H_2_CH_2_NH_2_), 116.5 (q, ^1^*JF* = 296.3 Hz, <u>C</u>F_3_COOH) 118.5 (pur C5), 142.9 (pur C8), 147.5 (pur C2), 149.3 (pur C4), 152.3 (pur C6), 158.8 (q, ^2^*JF* = 33.5 Hz, CF_3_<u>C</u>OOH).

### Ade-NBD

Synthesized according to ^7^ Redish-brown solid, yield 68 %, HPLC purity 99.9, ESI^+^-MS 342 (100, [M+H]^+^), ^1^H-NMR (500 MHz, DMSO-*d*_6_) δ (ppm): 3.92 (s, 2H, CH_2_C<u>H</u>_2_NHNBD), 4.46 (s, 2H, C<u>H</u>_2_CH_2_NHNBD), 6.44 (d, J = 8.3 Hz, 1H, NBD H6), 7.19 (s, 2H, NH_2_), 8.04 (s, 1H, pur C2), 8.09 (s, 1H, pur H8), 8.46 (d, J = 8.9 Hz, 1H, NBD H5), 9.48 (s, 1H, CH_2_CH_2_N<u>H</u>NBD). ^13^C-NMR (125 MHz, DMSO-*d*_6_) δ (ppm): 41.3 (<u>C</u>H_2_CH_2_NHNBD), 42.8 (CH_2_<u>C</u>H_2_NHNBD), 99.3 (NBD C6), 118.7 (pur C5), 121.3 (NBD), 137.7 (NBD C5), 141.0 (pur C8), 144.0 (NBD), 144.4 (NBD), 144.9 (NBD), 149.7 (pur C4), 152.3 (pur C2), 155.9 (pur C6).

